# Novel regulator Fpr2 controls immune homeostasis via CCL4-dependent neutrophils

**DOI:** 10.1101/2024.09.13.612885

**Authors:** Zeyu Sun, Song Mei, Shenglong Xu, Jiaxuan Yi, Shengwei Zhang, Yuhang Wang, Peng Liu, Hui Liu, Qingyu Lv, Wenhua Huang, Decong Kong, Qian Li, Yuhao Ren, Yongqiang Jiang, Peng Zhang, Hua Jiang

## Abstract

Polymorphonuclear leukocytes (PMNs) in *Fpr2-*deficient mice have significantly impaired bactericidal activity, yet the precise mechanism remains elusive. Our research unveiled the cellular heterogeneity of peripheral blood PMNs through single-cell RNA-sequencing (scRNA-seq) and identified two new neutrophil clusters related to bacterial infection, including a sterilization cluster named Neu_CSTA and a regulation cluster named Neu_CCL4. The lack of *Fpr2* leads to an immune regulation imbalance of Neu_CCL4 on other clusters in uninfected conditions, resulting in persistent instability of PMNs in peripheral blood, elevated inflammatory factors, and increased NOX2 expression. Furthermore, our findings corroborate that the regulatory function of Neu_CCL4 on PMNs immune balance is closely tied to its high expression of the anti-inflammatory ligand Anxa1. Regulon analysis inferred by scRNA-seq and ChIP-seq data suggested that a specific transcription factor Elf4, could modulate the expression of *Anxa1*, thereby facilitating its regulatory impact on other clusters. In the infection state, there was a noted increase in intercellular interactions across different clusters, leading to a general improvement in the bactericidal activity of PMNs. In summary, our research revealed distinct functional cellular sub-populations of PMNs, discovered the potential regulatory mechanisms, and elucidated the essential roles of Fpr2 in their ability to resist infection.

## Introduction

Neutrophils, as polymorphonuclear leukocytes within the phagocytic system, serve as the primary defense mechanism against invading pathogens and play a crucial role in mediating inflammation-induced injury (Silvestre-Roig, Hidalgo, & Soehnlein, 2016). It has been traditionally believed that neutrophils, being terminally differentiated cells, do not undergo further differentiation in peripheral blood and have a limited lifespan in this environment due to their inability to divide after differentiation. Consequently, limited research has been conducted on the phenotypic heterogeneity and functional diversity of PMN. Although the advancement of single-cell RNA sequencing(scRNA-seq) has revealed that PMN exhibits heterogeneity in peripheral blood (Beyrau, Bodkin, & Nourshargh, 2012; Xie et al., 2020), it remains uncertain whether there will be regulatory interactions among different clusters, and the molecular underpinnings of neutrophil heterogeneity are still ambiguous.

*Xie et al*. revealed the heterogeneity of neutrophils in the homeostasis and acute stages of infection for the first time through scRNA-seq and proposed that PMNs in peripheral blood are mainly mature PMNs, which have bactericidal-related functions (Xie et al., 2020). *Deerhake M. E. et al*. conducted a scRNA-seq analysis to investigate the heterogeneity of neutrophils during acute pulmonary Cryptococcus neoformans infection and posited the presence of oxidative stress characteristics (Ox-PMN) and distinct cytokine expression subsets. The varying functional subgroups of cells exhibited differing survival rates, indicating a close correlation between their functions and longevity (Deerhake, Reyes, Xu-Vanpala, & Shinohara, 2021). *Qi X. et al*. underscored the heterogeneity and functional diversity of neutrophils in the initial phase of severe burn patients, revealing notable variances in the dynamic equilibrium and predominant subsets of neutrophils in early burn patients as compared to individuals in good health (Qi et al., 2021). The aforementioned studies demonstrate the presence of distinct functional subsets of neutrophils in peripheral blood, highlighting the essential role of this diversity in host defense against infection.

FPR2 is a crucial chemotactic receptor expressed in many cells, with the highest levels found in myeloid cells. As a G protein-coupled receptor (GPCR), FPR2 interacts with a diverse array of ligands, thereby playing a multifaceted role in host immune responses (Corminboeuf & Leroy, 2015; He & Ye, 2017; Weiss & Kretschmer, 2018). Research has demonstrated the significance of FPR2 in host defense against *S. aureus* infection. However, our previous findings indicate that *S. aureus* induces vascular permeability through the activation of HBP release by FPR2 on PMNs (Kretschmer et al., 2010; Li et al., 2016). The absence of FPR2 leads to diminished host resistance to bloodstream infection with *GBS* (*Group B Streptococcus, GBS*) (Sun et al., 2021) yet the activation of FPR2 during *S suis* infection intraperitoneally can lead to severe inflammatory response and an increase in mortality rates (Ni et al., 2023). Studies of *L. monocytogenes*, *E. coli*, and *S. pneumoniae* infections have shown similar phenomena (Liu et al., 2012; Machado et al., 2020; M. Zhang et al., 2020). These findings indicate that FPR2 plays a significant role in modulating the inflammatory response, with a complex underlying mechanism. This receptor is likely a crucial target for both host and pathogenic bacteria in the context of infection, necessitating further investigation.

Currently, the majority of functional investigations of FPR2 are centered on its role in various diseases, with a limited understanding of its involvement in immune homeostasis. Some researchers have demonstrated the significance of constitutive *FPR2* expression in the maintenance of immune homeostasis (Dufton et al., 2010). In the absence of exogenous agonists, transgenic mice with bone marrow cells overexpressing human *FPR2/ALX* showed reduced neutrophil infiltration in a yeast polysaccharide-induced peritonitis model (Devchand et al., 2003). Macrophages in *Fpr2*-deficient mice had reduced responses to pro-inflammatory ligands fMLP, SAA, and anti-inflammatory ligands annexin A1 (ANXA1) and LXA4, but mice showed a more pronounced inflammatory response overall (Dufton et al., 2010). LXA4 can enhance the activity of *FPR2/ALX* promoter and the overexpression of *FPR2/ALX* in transgenic mice is related to the decrease of inflammation (Simiele et al., 2012). In the case of *FPR2* deletion or reduced expression, the host showed varying degrees of anti-inflammatory dysregulation. For example, *FPR2* promoter polymorphism rs11666254 was significantly associated with sepsis in trauma patients. Also, a remarkable correlation was identified between reduced *FPR2* expression and elevated TNF-α levels in peripheral blood leukocytes stimulated by LPS (H. Zhang et al., 2017). Mutations C126W and F110S in the second intracellular loop and third transmembrane region of FPR2 lead to a partial or complete defect in G protein-coupled receptors signaling, particularly in adolescent periodontitis patients rather than in adults (Seifert & Wenzel-Seifert, 2001). Our previous studies have also indicated that *Fpr2*-deficient mice exhibited increased inflammation and compromised host resistance to pathogens with impaired PMN chemotaxis and bactericidal capabilities (Sun et al., 2021). However, the specific mechanism is not very clear. In this study, the regulation of FPR2 on PMNs’ function and immune balance was clarified through scRNA-seq and experimental validation.

## Results

### The lack of *Fpr2* increases the production of NOX2 and inflammatory factors, while reduces ROS expression

Previous studies have indicated an association between FPR2 and host immune homeostasis. Here it was also observed that *Fpr2*-deficient mice exhibited a certain degree of immune disorder. The alteration in PMNs was examined through scRNA-seq and validated using a series of biological assays (**Fig.1a**). Analysis of peripheral blood samples from *Fpr2*-KO mice (KO) and wide type (WT) mice via ELISA reveals the imbalance of inflammatory factors, characterized by a notable elevation of pro-inflammatory cytokines IL-1 β, IL-6 and IL-17, as well as the anti-inflammatory cytokine IL-4 (**Fig.1b-e**). Furthermore, PMN function disruption was investigated in KO mice. The expression of NADPH-oxidase 2 (NOX2) genes *Cybb*, *Cyba*, secondary secretory granule-related genes *Itgam*, *Mmp25*, and ROS-related genes *Mpo* and *Hsp90* was detected by qPCR. The results showed that in the steady-state, the ROS generation and granule secretion of PMN in *Fpr2*-KO mice decreased, while NOX expression increased (**Fig.1f-h**). ELISA was employed to assess the expression level of NOX2. Aligning with the gene expression data, the content of NOX2 in the PMN of KO mice exhibited a marked increase (**Fig.1i**).

To further validate these findings, peripheral blood neutrophils from healthy human donors were collected and exposed to the receptor antagonist WRW4. The results demonstrated a significant increase in the expression of *Cyba* (**Fig.1j**). Interestingly, unlike in murine models, the expression of *Itgam* was also significantly up-regulated, while the expression of *MPO* was notably decreased and no significant difference of *Mmp25* was observed (**Fig.1k-m**). These results indicated that degranulation disruption, ROS generation, and NADPH elevation occurred in the PMNs of *Fpr2*-KO mice.

### Single-cell transcriptomic analysis deciphers cellular changes of neutrophils related to *Fpr2* deficiency

To further clarify the regulatory mechanism of FPR2 in immune homeostasis and the bactericidal function of PMNs, we collected peripheral blood immune cells from *Fpr2*- KO or WT in resting or *GBS* infected condition and performed scRNA-seq. After stringent quality control, graph-based clustering and canonical markers annotation revealed 4 myeloid immune cell populations, including dendritic cells (n=256), macrophages (n=1,526), monocytes (n=2,441) and neutrophils (n=17,363)(**Fig. 2a,b, Supplementary Table S1**). Neutrophils was the major myeloid immune cell type, especially in *GBS* infection (**Fig. 2c**). In resting condition, the proportion of neutrophils in peripheral blood elevated when *Fpr2* was knocked out while no other myeloid cell types had statistical changes. On the other hand, *GBS* infection caused a decrease in both the proportion and absolute number of macrophages, monocytes, and DCs, while neutrophils increased remarkably (**Fig. 2d**).

**Fig. 1.**
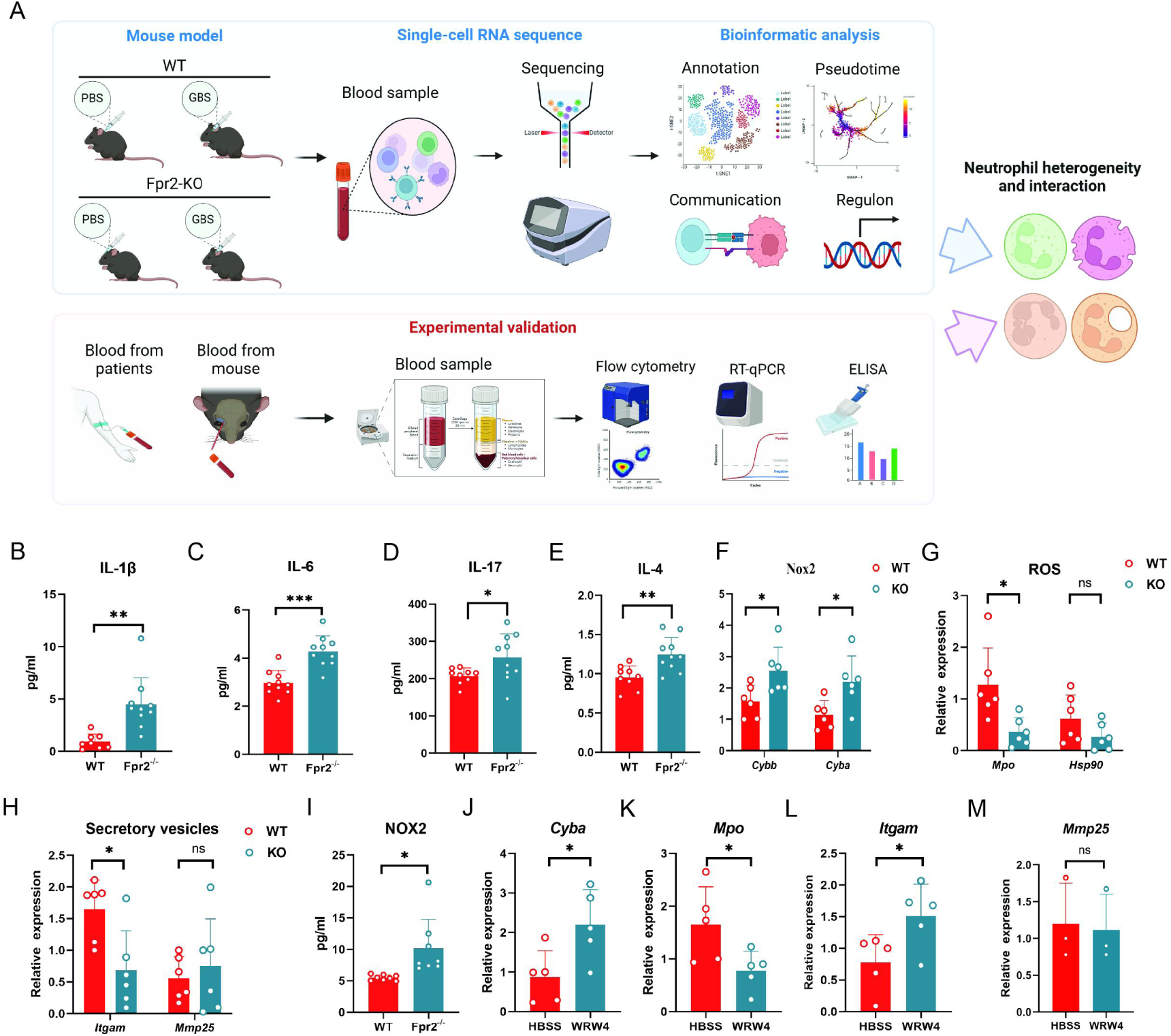
The absence of *Fpr2* results in dysregulation of inflammatory cytokine levels and impaired PMN function in homeostasis. **A** The framework about our research. **B-E** The deletion of *Fpr2* results in dysregulation of IL-1β (**B**), IL-6 (**C**), IL-17 (**D**) and IL-4 (**E**) expression in uninfection state. Data are representative of two independent experiments with n = 10 mice per group. Data are shown as *t test*. *\*p* < 0.05, \*\**p* <0.01. ns, denotes not significant. **F-H** The absence of *Fpr2* results in impaired neutrophil NADPH (**F**), ROS (**G**) and Secretory vesicles (**H**) in uninfection state. Data are representative of two independent experiments with n=6 mice per group. Data are shown as t test. *\*p* < 0.05, ns, denotes not significant. **I** Increased NOX2 expression in PMN of *Fpr2*-KO mice by ELISA. Data are representative of three independent experiments with n=8 mice per group. Data are shown as *t test*. *\*p* < 0.05. **J-M** WRW4 treatment causes *Cyba* (**J**), *Mpo* (**K**), *Itgam* (**L**) and *Mmp25* (**M**) dysfunction in human peripheral blood PMN. Data are representative of two independent experiments with n=5 mice per group. Data are shown as *t test*. *\*p* < 0.05, ns, denotes not significant.

**Fig. 2.**
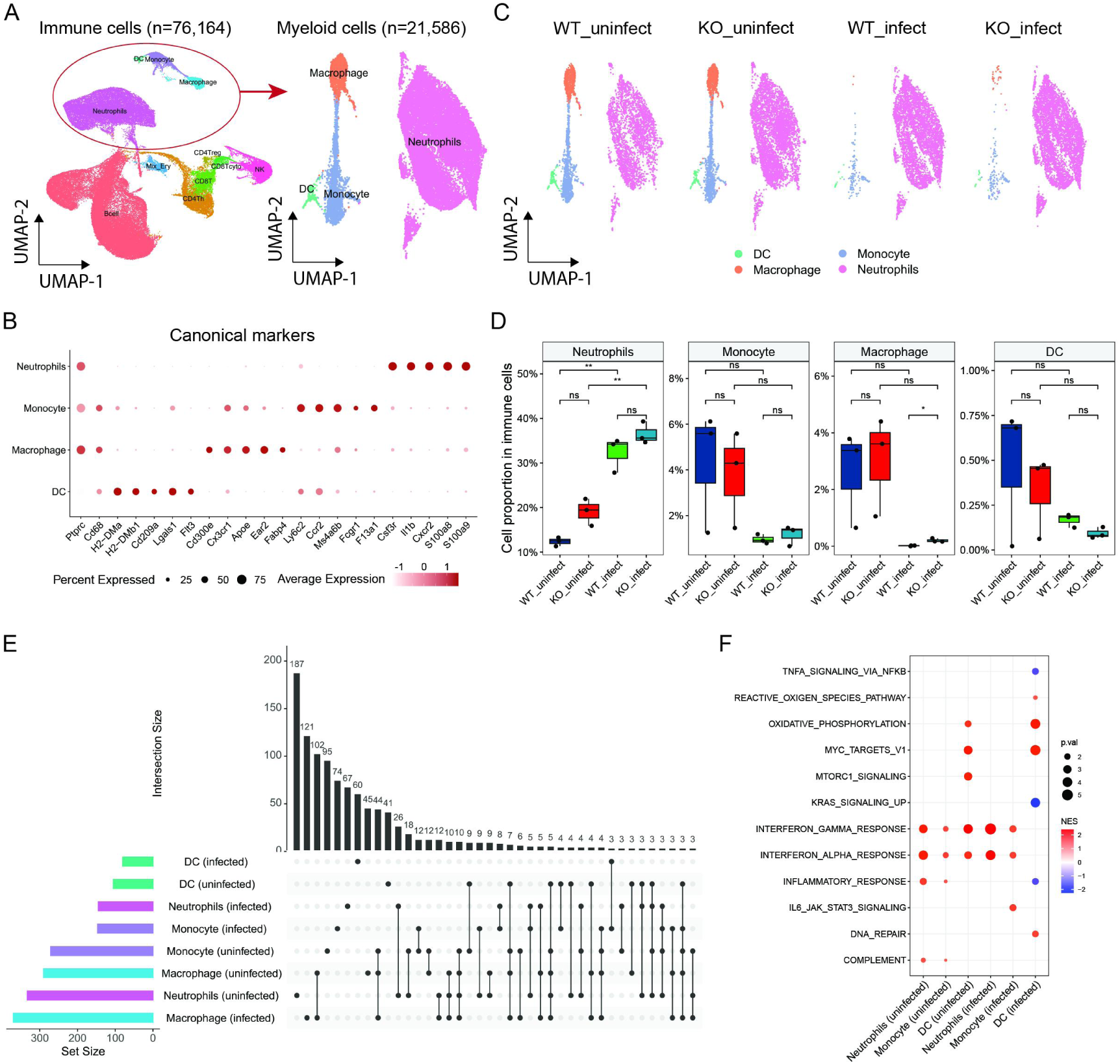
scRNAseq profile of immune cells in *Fpr2* knock-out and *GBS* infection. **A** The UMAP plot of single cell clusters. **B** Dot plot of corresponding canonical markers of each cell cluster. **C** The UMAP plot of each cluster, divided in WT_uninfect(n=3), KO_uninfect(n=3), WT_infect(n=3) and KO_infect(n=3) groups. **D** Boxplot of each cell cluster’s proportion in myeloid immune cells, divided in WT_uninfect, KO_uninfect, WT_infect and KO_infect groups. Statistical analysis was performed by stat_compare_means function in ggpubr package. **E** Counts of DEGs between KO and WT (left bar plot) and the overlap DEGs number between cell clusters (top bar plot and dot plot with lines) under uninfect or infect condition. **F** Heatmap of GSEA results based on DEGs calculated in (**e**), with Hallmark gene sets as the reference. Data are shown as *t test. *p* < 0.05, \*\**p* <0.01. ns, denotes not significant.

To explore the function of immune cells influenced by Fpr2 in case of resting or infection, we also calculated differential expressed genes (DEGs) between *Fpr2*-KO and WT. Compared with resting condition, *Fpr2* deficiency induced different gene expression patterns of myeloid cells under *GBS* infection (**Fig. 2e**). Furthermore, more DEGs were induced in neutrophils by *Fpr2* knock out in resting condition than infection (187 DEGs vs. 67 DEGs, **Fig. 2e**). Through GSEA analysis, IFN-α/γ response pathways in neutrophils and monocytes were main inflammatory processes up-regulated by *Fpr2* knock out (**Fig. 2f**). After *GBS* infection, gene expression of macrophages was less affected by *Fpr2* deficiency (**Fig. 2e**). IFN-α/γ response pathways were also elevated in neutrophils and monocytes, but TNFα signaling via NF-κB went down in DCs (**Fig. 2f**). These changes indicated the lack of *Fpr2* could affect the immune response of PMN both with and without infection and indicated neutrophils was the most likely participant in *GBS* infection.

### Neu_CSTA and Neu_CCL4 were the main neutrophil subtypes affected by Fpr2

The dramatic variation of both cell proportion and gene expression of neutrophils inspired us to explore the regulatory function of Fpr2 in *GBS* infection, so we further re-clustered the neutrophil population into 8 subtypes based on their canonical markers (**Fig.3a,b**, **Supplementary Table S2**). For a better known of each subtype’s function, we enriched the biological processes in Gene Ontology based on genes specifically expressed in each subtype (**Fig. 3c**). The major neutrophil cluster (which was named Neu_classic), was a classical neutrophil subcluster with no specific functional markers. Activated neutrophil(Neu_mt-ND1), with mitochondrial genes highly expressed, showed the function of IL-8, IL-6, and TNF production. Neu_CCL4 could respond to IFN-γ and produce chemokines like CCL4 and CCL3, which may regulate neutrophil recruitment and inflammatory function (Kolaczkowska & Kubes, 2013). Neu_CD177 was a subtype with elevated glucose 6-phosphate metabolic process and related to cell activation. Neu_CSTA highly expressed *Cstdc4* and *Stfa2l1*, which are orthologous to human CSTA and indicated to be related to neutrophil degranulation and leukocyte-mediated immunity (Ahmed et al., 2023). Neu_CTLA2A participated in homotypic cell-cell adhesion. Neu_ISG15 was the main subtype in response to IFN-α/β and played roles in antigen processing and presentation. Neu_NINJ1 highly expressed genes related to cell differentiation and adhesion (**Fig. 3c**).

**Fig. 3.**
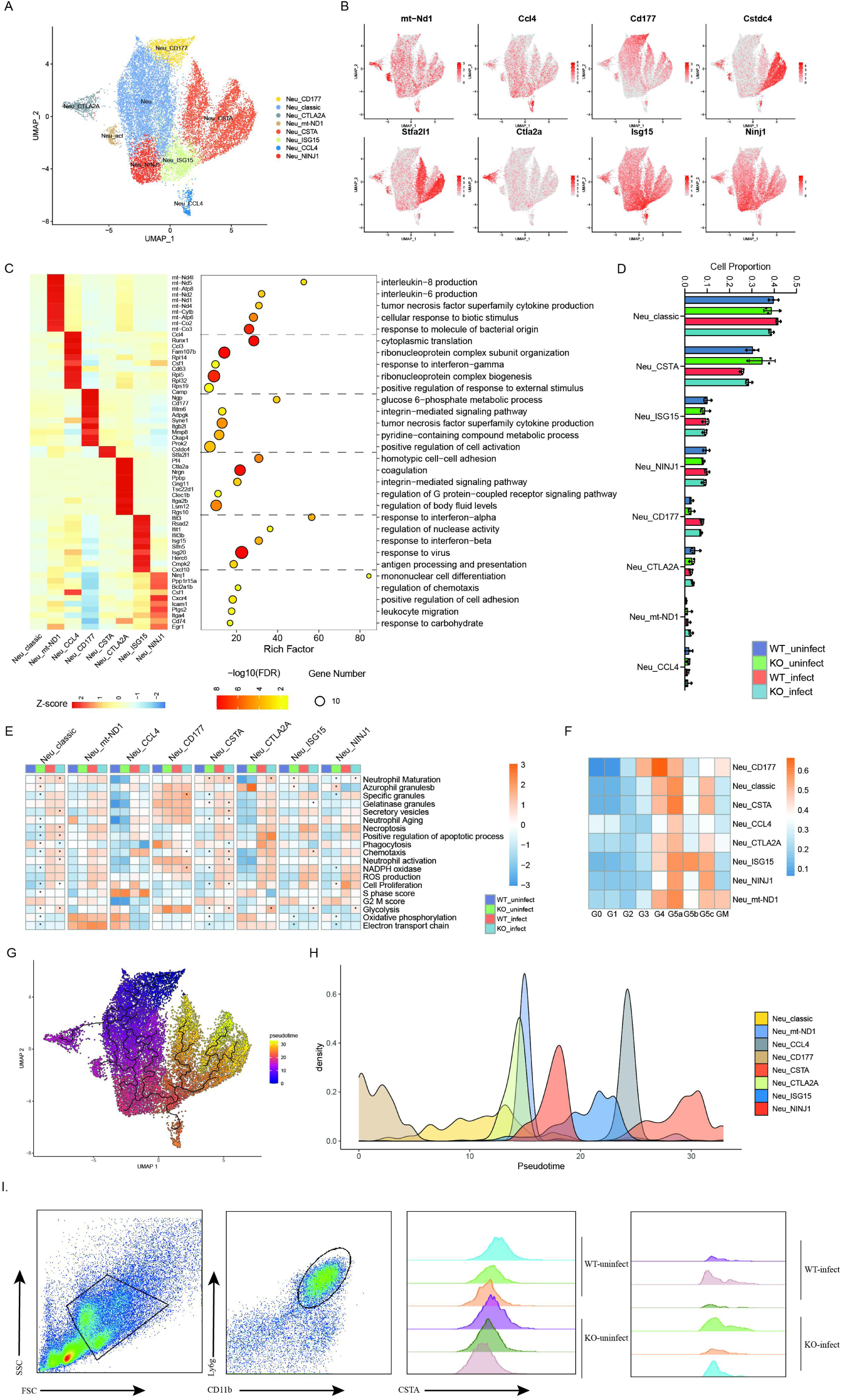
Neu_classic and Neu_CSTA were main subtypes in response to *GBS* infection and influenced by *Fpr2* knock-out. **A** The UMAP plot of neutrophil subtypes. **B** The expression of canonical markers in each neutrophil subtype. **C** Heatmap of highly expressed and specific genes in each neutrophil subtype (left) and the enrichment results of these genes (right). **D** Boxplot of each neutrophil subtype’s proportion in all neutrophils, divided in WT_uninfect, KO_uninfect, WT_infect and KO_infect groups. **E** Heatmap of GSVA scores of crucial functions identified in previous research related to neutrophils. **F** Heatmap of GSVA scores of cell cycle-related gene sets. **G** The UMAP plot of neutrophils, colored by pseudo-time and cell trajectory was shown in black lines. **H** Density plot of the distribution of each neutrophil subtypes along the pseudo-time. **I** Flow cytometry detection of CSTA in peripheral blood of WT_uninfect, KO_uninfect, WT_infect and KO_infect groups. Data are representative of two independent experiments with n=3 mice per group. Data are shown as *t te*st. *\*p*< 0.05, \*\**p*<0.01. ns, denotes not significant.

These findings corroborated the functional heterogeneity of PMNs in peripheral blood. However, the proportion of each neutrophil subtype just showed slight variation between infected and resting conditions, or between *Fpr2*-KO and WT. *GBS* infection could increase the proportion of Neu_CD177 no matter whether *Fpr2* was knocked out. However, *Fpr2* knock out slightly decreased the proportion of Neu_classic and faintly increased the proportion of Neu_CSTA (**Fig. 3d**). Then we further investigated some crucial functions identified in previous research related to neutrophils. *Fpr2* deficiency remarkably affected the function of Neu_classic and Neu_CSTA subtypes in both uninfected and infected conditions, *i.e.* maturation, chemotaxis, proliferation, glycolysis (**Fig. 3e**). Meanwhile, *Fpr2* deficiency could also inhibit the aging of these two subtypes (**Fig. 3e**). Since the main functions of PMN subsets in peripheral blood are different, we attempted to identify the developmental trajectory of neutrophils. Firstly, we calculated the score of neutrophils in different cell cycle phases from previous research (Xie et al., 2020). Neu_CD177 was mainly similar to neutrophils in G4 and partially like neutrophils in G3, while other subtypes were likely in G5a/b/c stage (**Fig. 3f**). The increase in CD177 during the infection mentioned above may be because infection promotes the accelerated release of PMN from the bone marrow, which has been confirmed by other studies (Xie et al., 2020). Therefore, we assumed Neu_CD177 was the subtype in the early developmental stage and performed the pseudo-time analysis. As shown in **Fig. 3g** and **3h**, Neu_CTLA2A (cell adhesion), Neu_mt-ND1 (cytokines production), Neu_CCL4 (chemokines production) and Neu_CSTA (degranulation) were different endpoints of cell trajectories which again confirms the development track of PMN discharged from bone marrow to different functions in peripheral blood again and to some extent explained the heterogeneity of neutrophil subtypes in mouse. Each subtype has distinctive gene expression patterns and functions, among which Neu_CSTA was the main functional subcluster in response to *GBS* infection. Fpr2 dramatically influenced the function of Neu_CSTA and may prompt more neutrophils in the Neu_classic subtype to develop into Neu_CSTA in *GBS*-infected condition. Given that CSTA is the primary bactericidal functional group, our observation focused on the quantity of CSTA^+^ cells in *Fpr2*-KO mouse neutrophils. It revealed that the levels of CSTA^+^ cells in peripheral blood remained consistent before and after *Fpr2* knockout with or without infection (**Fig. 3i**).

### Neu_CSTA was the most influential sub-population of PMN by *Fpr2* deficiency

Because Neu_CSTA is the most affected function cluster with *Fpr2* deletion, to clarify the function of Neu_CSTA and their alteration in *Fpr2* deficiency and *GBS* infection, we further clustered it into three populations based on their specific markers, including Cstdc4^+^Stfa2l1^-^ (Neu_Cstdc4), Cstdc4^-^Stfa2l1^+^ (Neu_Stfa2l1) and Cstdc4^+^Stfa2l1^+^ (Neu_Double Positive, Neu_DP) (**Fig. 4a,b**). Then we analyzed the DEGs between *Fpr2*-KO and WT in resting or *GBS*-infected conditions, respectively. In resting condition, most DEGs were shared among three populations. Neu_DP and Neu_Stfa2l1 had more unique or shared DEGs (**Fig. 4c**). KEGG pathways were enriched to identify the influence of Fpr2 on three populations. Pathways related to microbial or parasitic infection were changed by *Fpr2* knock-out. For Neu_Stfa2l1, FcεRI signaling pathway, FcγR-mediated phagocytosis, and TLR signaling pathway were altered. C-type lectin receptor signaling pathway and Rap1 signaling pathway were unique in Neu_DP and Neu_Cstdc4, respectively (**Fig. 4d**). In *GBS* infection, DEGs shared by three populations were less than normal conditions. Unlike uninfected situation, Neu_DP and Neu_Stfa2l1 also had numerous unique but less shared DEGs (**Fig. 4e**). KEGG pathway enrichment analysis indicated *Fpr2* knock-out also affected response to microbial infection (including RIG-I-like receptor, Toll-like receptor and NOD-like receptor signaling pathways) in all populations. For Neu_Stfa2l1, VEGF and HIF-1 signaling pathways were changed, which was not seen in the uninfected situation. Antigen processing and presentation, phagosome, and endocytosis showed a difference in Neu_DP and NF-κB signaling pathway altered in Neu_Cstdc4 (**Fig. 4f**). These results together demonstrated that sub-populations of Neu_CSTA showed different immunologic functions, especially in infected situation, of which Neu_Stfa2l1 was most likely affected by Fpr2.

**Fig. 4.**
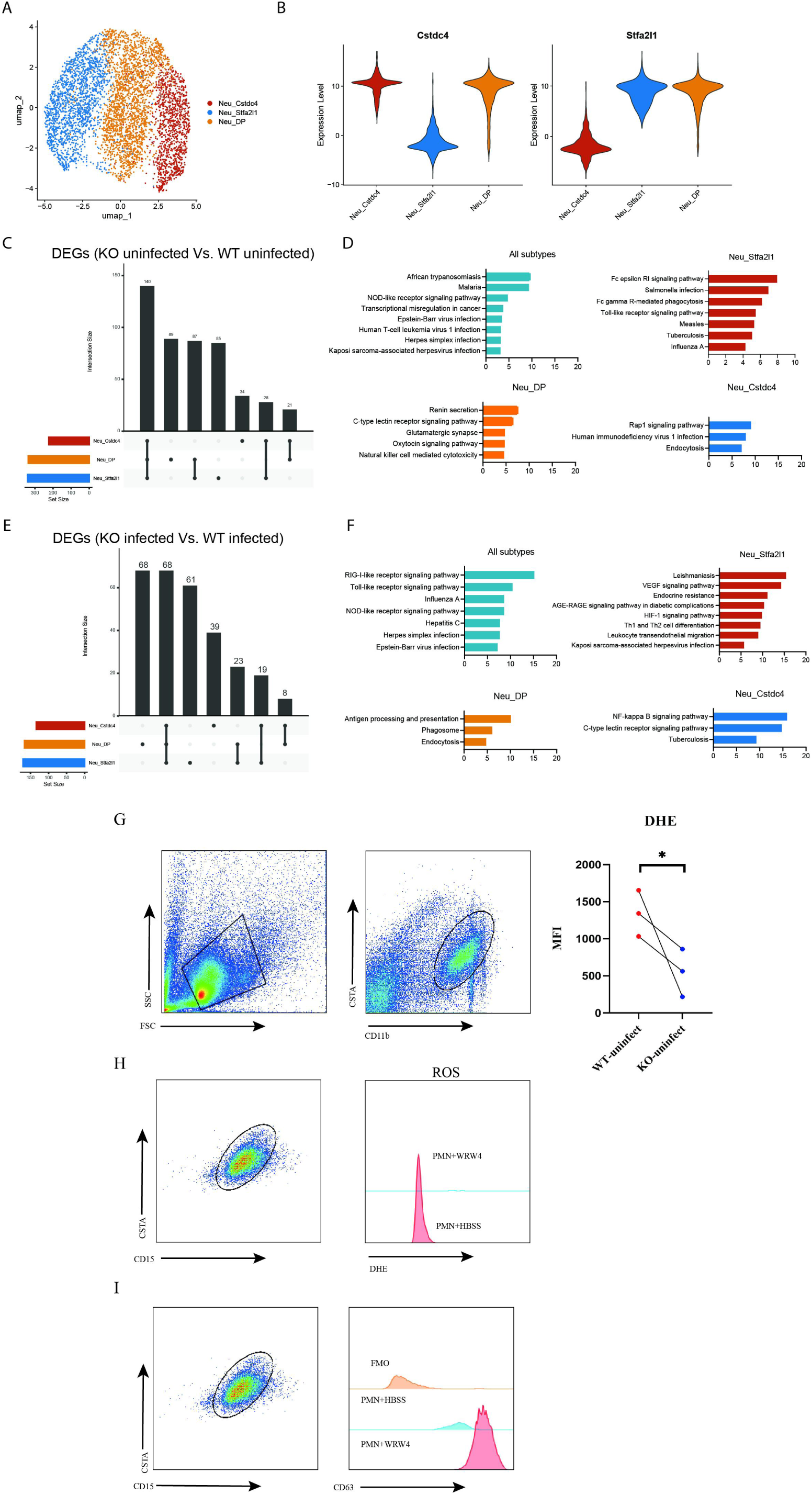
Neu_Stfa2l1 was the main functional neutrophil population affected by Fpr2. **A** The UMAP plot of three Neu_CSTA populations. **B** Violin plots of specific markers expressed in each Neu_CSTA populations. **C** The number of DEGs between WT_uninfect and KO_uninfect in each Neu_CSTA populations or shared by these populations. **D** Enrichment results of KEGG pathways based on DEGs shared by three populations or unique in each population, corresponding to (**c**). **E** The number of DEGs between WT_infect and KO-infect in each Neu_CSTA populations or shared by these populations. **F** Enrichment results of KEGG pathways based on DEGs shared by three populations or unique in each population, corresponding to (**E**). **G-H** Flow cytometry detection of the changes in ROS expression in mouse (**G**) and human (**H**) CSTA clusters with *FPR2* deficiency. **I** Flow cytometry detection of the changes in CD63 expression in human CSTA cluster with WRW4 treatment. Data are representative of two independent experiments with n=3 mice per group. Data are shown as *t te*st. *\*p* < 0.05, \*\**p* <0.01. ns, denotes not significant.

Given the significant role of CSTA as a functional subgroup, our study sought to investigate the potential regulatory role of Fpr2 on ROS production and granule secretion within this subgroup. Our findings indicate that the lack of *Fpr2* leads to a notable decrease in ROS production in the uninfected state (**Fig. 4g,h**) and a significant increase in the granules marker CD63 (**Fig. 4i**). These results suggest that Fpr2 might be a key player in modulating the bactericidal activity of PMN function cluster.

### Neu_CCL4 was the main regulatory sub-population in neutrophil homeostasis during bacterial infection

Given the minimal impact of *Fpr2* knockout on the distribution of CSTA subsets, yet its significant influence on their functionality and PMN in peripheral blood is divided into many clusters and the clusters are not independent, we speculate whether Fpr2 may affect PMN through inter-subgroup interaction. To verify this hypothesis, we performed a cell-cell communication analysis. In resting condition, Neu_CCL4 had the highest outgoing interaction strength in all neutrophil subtypes, while Neu_CD177, Neu_classic and Neu_CSTA received the strongest incoming signals **(Fig.5a**). We further discovered CCL pathway was a universal signaling axis from Neu_CCL4 to other neutrophil subtypes (**Fig. 5b**). CCL pathway decreased in *GBS* infection and further declined after *Fpr2* knocked out. In the context of infection, peripheral blood PMN clusters exhibit enhanced activation of bactericidal activity and a marked augmentation in inter-subgroup interactions to efficiently eradicate pathogenic agents (**Fig. 5b,c**).

**Fig. 5.**
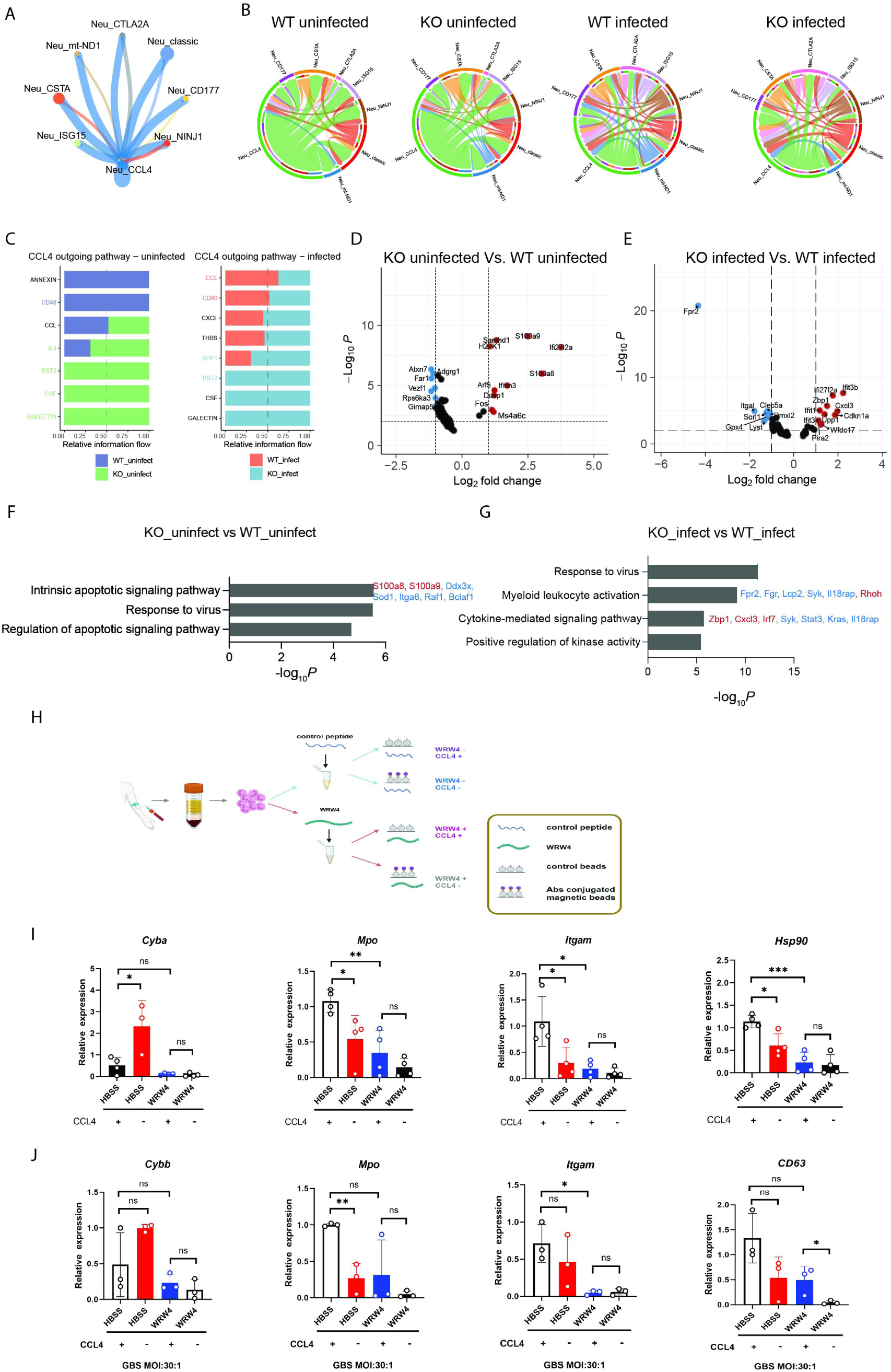
Cell-cell communications between neutrophil subtypes. **A** The network of outgoing interactions of Neu_CCL4. **B** The chord plots of CCL pathway between neutrophil subtypes in WT_uninfect, KO_uninfect, WT_infect and KO_infect groups. **C** The difference of outgoing pathway of Neu_CCL4 between WT_uninfect and KO_uninfect (left) and between WT_infect and KO_infect (right). **D,E** Volcano plots of DEGs between WT_uninfect and KO-uninfect (**D**) and WT_infect and KO_infect (**E**) in Neu_CCL4. **F,G** Enrichment results of KEGG pathways in Neu_CCL4 based on DEGs between WT_uninfect and KO_uninfect (**F**) or WT_infect and KO_infect (**G**). **H** Schematic representation of the processing of Neu_CCL4 and WRW4. **I-J** Gene expression in human peripheral blood PMNs following treatment with WRW4 or deletion of Neu_CCL4, in uninfection (**I**) or in infection (**J**). Data are representative of two independent experiments with n = 3-4 mice per group. Data are shown as *t te*st. *\*p* < 0.05, \*\**p* <0.01. ns, denotes not significant.

The chemokine ligand *CCL4*, which binds to CCR5, was highly expressed in the Neu_CCL4 subtype (**Fig. 3b**). In normal condition, *Fpr2* knock-out up-regulated *S100a8* and *S100a9* in Neu_CCL4, which play prominent roles in neutrophil-related inflammatory processes and response (**Fig. 5d,e**). However, more genes related to the intrinsic apoptosis signaling pathway except for *S100a8* and *S100a9* were down-regulated by *Fpr2* knock-out in resting condition (**Fig. 5f**). After *GBS* infection, myeloid leukocyte activation process and cytokine-mediated signaling pathway were disrupted by *Fpr2* knock-out, which indicated the function of Neu_CCL4 was also governed more by Fpr2 under infected condition (**Fig. 5g**).

From **Fig. 3E**, we found that Neu_CCL4 had relatively low levels of degranulation, secretory vesicles, and phagocytosis. Based on the function analysis in **Fig.3C** and **3E**, Neu_CCL4 was not a typical bactericidal subcluster. And in consideration of the chemokine secretion function of Neu_CCL4, we assumed it could function as a regulatory subtype. To confirm the potential regulatory impact of Neu_CCL4 on other clusters and its relationship with Fpr2, we treated PMNs from healthy human blood samples with WRW4 and utilized magnetic bead sorting to simulate the effects of two distinct states: one with Neu_CCL4 present and one without (**Fig. 5h**). The findings indicate that the lack of Neu_CCL4 cluster resulted in a significant increase in *Cyba* expression and a decrease in *Mpo*, *Itgam* and *Hsp90* expression under rest conditions. Treatment of PMN with WRW4, either alone or in combination with Neu_CCL4 deletion, resulted in a notable decrease in the expression levels of *Cyba*, *Mpo*, *Itgam*, and *Hsp90*, and there was no significant difference observed between the groups(**Fig. 5i**). These findings suggest a close association between Neu_CCL4 regulatory and Fpr2 in the resting condition.

In cases of infection, deletion of Neu_CCL4 resulted in a decrease in the expression of *Mpo*, *Itgam*, and *CD6*3, but not *Cybb*. There is a notable disparity in the extent of reduction in *Mpo* expression (**Fig. 5j**). This indicates that without Neu_CCL4, the ability of PMNs to effectively eliminate bacteria may be compromised to a certain extent. Subsequently, it was observed that the administration of the Fpr2 antagonist WRW4 led to a notable decrease in the expression levels of *Mpo*, *Itgam*, *CD63*, and *Cybb*, regardless of the presence of Neu_CCL4. Specifically, the expression of *CD63* was significantly reduced to nearly undetectable levels (**Fig. 5j**). These results highlighted the significance of Neu_CCL4 as a regulatory factor of PMNs both during homeostasis and in infection.

### The expression of Anxa1 was modulated by Elf4, which acts on Fpr2 to control the equilibrium of PMN clusters

Furthermore, we want to elucidate the relationship between the regulation function played by Neu_CCL4 and Fpr2. As Neu_CSTA was the main functional subtype response to *GBS* infection and regulated by Fpr2, we focused on the relative interaction flow from Neu_CCL4 to Neu_CSTA. In resting condition, interactions including Lgals9-Ighm/Cd45/Cd44, Csf1-Csf1r, and Bst2-Pira2 were up-regulated, while Cd48-Cd244a and Anxa1-Fpr2 were down-regulated after knocking out *Fpr2* (**Fig. 6a**). ELISA confirmed the significant decrease in the concentration of Anxa1 in the peripheral blood of *Fpr2*-KO mice (**Fig. 6e**). On the other hand, after *GBS* infection, Spp1-Cd44 and Cxcl3-Cxcr2 was elevated and Spp1-(Itga4+Itgb1) was decreased by *Fpr2* knock-out (**Fig. 6a**).

**Fig. 6.**
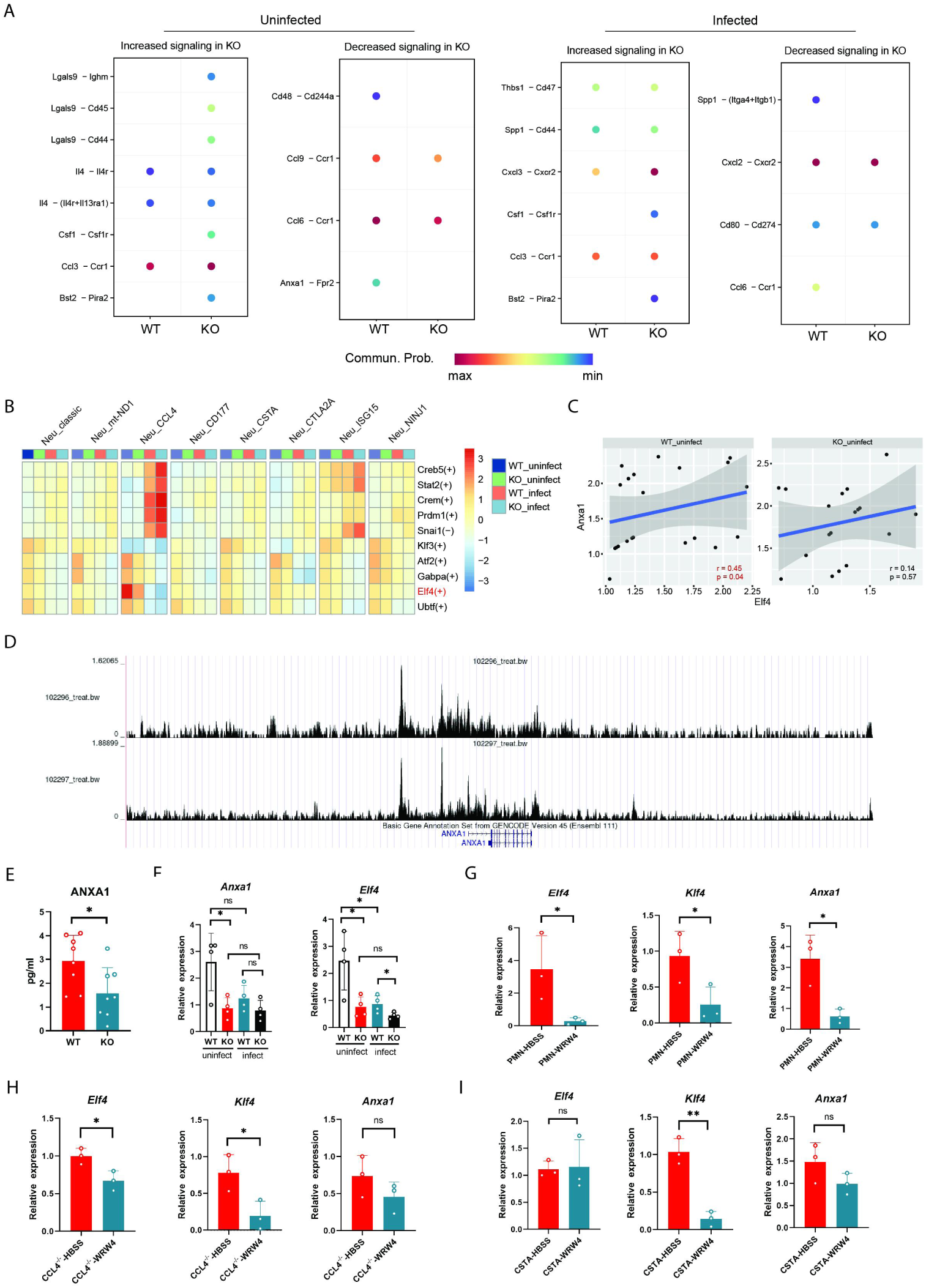
Transcription factor Elf4 regulates the modulatory function of Neu_CCL4. **A** The heatmap of the ligand-receptor pairs between Neu_CCL4 and Neu_CSTA in WT_uninfect, KO-uninfect, WT_infect and KO_infect groups. **B** Heatmap of regulon specificity score of each neutrophil subtype, divided in WT_uninfect, KO-uninfect, WT_infect and KO_infect groups. **C** The Spearman correlation between the expression of *Elf4* and *Anxa1* in Neu_CCL4, divided in WT_uninfect and KO-uninfect groups. **D** The peak chart of the potential Elf4-binding cite on the upstream of *Anxa1*. **E** ANXA1 concentration in mice detected by ELISA. **F** Expression of *Anxa1* and *Elf4* in mice in infection and uninfection state. **G-I** Expression levels of *Elf4*, *Klf4* and *Anxa1* in human peripheral blood PMN following WRW4 treatment (**G**), deletion of Neu_CCL4 and WRW4 treatment (**H**) and in Neu_CSTA cluster(**I**). Data are representative of two independent experiments with n=3-4 mice per group. Data are shown as *t te*st. *\*p* < 0.05, \*\**p* <0.01. ns, denotes not significant.

In order to find the reason for the formation of Neu_CCL4 phenotype and its relationship to Fpr2 regulation, we applied SCENIC analysis to identify the subtype-specific transcription factors(TFs). We found that Elf4 regulon was specifically expressed in Neu_CCL4 cluster (**Fig. 6b**). Also, Elf4 was the only Neu_CCL4-specific TF in resting condition (**Fig. 6b**). In the absence of infection, a correlation analysis revealed a statistically significant positive relationship between the expression levels of the transcriptional regulator Elf4 and *Anxa1* (**Fig. 6c**), which is a known anti-inflammatory regulator of Fpr2 as documented in existing literature (Perretti & D’Acquisto, 2009). RT-qPCR demonstrated the down-regulation of *Elf4* and *Anxa1* expression in response to *Fpr2* deletion under rest condition. Following *GBS* infection, a notable decrease in the expression of these two genes was observed, irrespective of the presence or absence of *Fpr2* (**Fig. 6f**). Previous studies did not find a direct link between Elf4 and Anxa1. Through analysis of the Elf4 ChIP-seq data from the Cistrome Data Browser (Consortium, 2012), we observed that Elf4 possesses a binding region upstream of *Anxa1* (**Fig. 6d**), suggesting a potential regulatory role of Elf4 on *Anxa1.* Otherwise, we also discovered several studies have indicated that Elf4 is capable of modulating the expression of *Klf4* (Yamada, Park, Mamonkin, & Lacorazza, 2009), while Klf4 in turn can directly influence the expression of *Anxa1* (Chen et al., 2023).

To clarify the association between Elf4 and Neu_CCL4, Neu_CSTA, and Fpr2, human PMNs were collected and subjected to depletion of Neu_CCL4, separation of Neu_CSTA, or treatment with WRW4. The findings indicated that in resting condition, the absence of *Fpr2* resulted in a notable decrease in the expression of *Elf4*, *Klf4*, and *Anxa1* concurrently (**Fig. 6g**). Upon deletion of the Neu_CCL4 cluster, the absence of *Fpr2* led to a significant reduction in *Elf4* and *Klf4*. Notably, it was evident that the overall expression of *Elf4* significantly decreased following the deletion of Neu_CCL4. Deletion of *Fpr2* did not result in a notable decrease in *Anxa1* expression in the absence of the Neu_CCL4 cluster. However, it was evident that the deletion of Neu_CCL4 itself significantly reduced *Anxa1* expression levels (**Fig. 6h**). These results indicate that Neu_CCL4 and its expression of *Fpr2* plays a significant role in the regulation of *Anxa1* expression.

As Neu_CSTA is a bactericidal function group, the expression levels of *Elf4*, *Klf4*, and *Anxa1* were examined within Neu_CSTA. The findings reveal that Neu_CSTA exhibits low expression levels of *Elf4* and *Anxa1* regardless of the presence of *Fpr2*. However, the absence of *Fpr2* results in a significant decrease in *Klf4* expression by Neu_CSTA(**Fig. 6i**). It is evident that Neu_CCL4 is the primary cluster expressing *Elf4* and *Anxa*1, with the expression of *Fpr2* playing a crucial role in regulating the expression of *Elf4*, *Klf4*, and *Anxa1*. However, the expression of *Klf4* is not exclusive to Neu_CCL4.

Based on our analysis, it is proposed that Neu_CCL4 may play a regulatory role in maintaining cellular cluster homeostasis by upregulating *Elf4* expression, which subsequently controls *Anxa1* expression. Furthermore, Fpr2 is involved in positively regulating both *Elf4* and *Anxa1* expression and also acts as a target of Anxa1. The transcription factor Klf4 may serve as a potential link between *Elf4* and *Anxa1*, or it is possible that Elf4 directly influences the expression of *Anxa1*, although further experiment is required to validate this hypothesis (**Fig.7**).

**Fig. 7.**
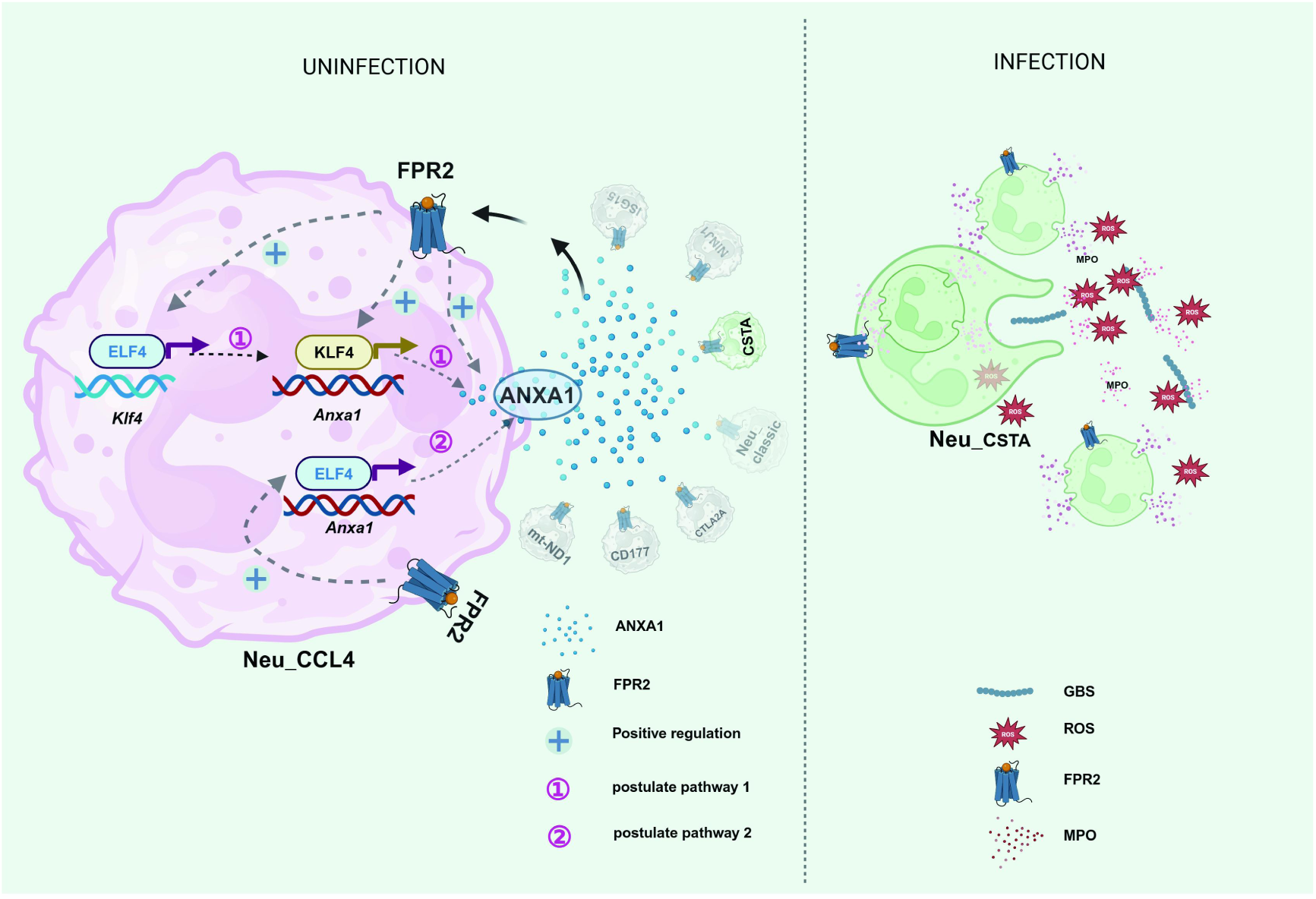
The schematic diagram illustrates the role of FPR2 in modulating the immune balance of PMNs through the regulation of ELF4*-*Anxa1 in Neu_CCL4 cluster.

## Discussion

This study conducted single-cell RNA sequencing analysis to investigate the regulatory effect of Fpr2 on immune balance and sterilization of PMN. The results corroborated the heterogeneity among PMNs in peripheral blood and clarified the potential regulatory mechanism within clusters. A novel bactericidal functional subgroup, Neu_CSTA, as well as an immune balance regulation subgroup, Neu_CCL4, were identified. In the steady state, Neu_CCL4 maintains immune balance and exerts regulatory effects on other subgroups through the ANXA1-FPR2 axis. The expression of *Anxa1* might be associated with the Neu_CCL4-specific regulon ELF4 and possibly the transcription factor KLF4. During bacterial infection, the regulatory influence of Neu_CCL4 decreased, instead intergroup interactions increased and overall bactericidal activity enhanced.

Prior research has established the significance of Fpr2 in modulating the inflammatory equilibrium of the host. Our study revealed that *Fpr2*-KO mice exhibited dysregulated expression of inflammatory cytokines in peripheral blood under basal conditions, along with significantly elevated expression of NADPH subunits *Cyba* and *Cybb* in PMN. Following treatment with WRW4, *Cyba* expression in human PMN was notably increased. These findings suggested that the absence of functional FPR2 in PMN disrupted immune homeostasis. Multiple studies have verified the heterogeneity of the PMN population, with distinct subsets exhibiting varied functions (Xie et al., 2020). This investigation revealed the presence of diverse PMN subpopulations in peripheral blood and highlighted the interplay between these subsets in the bactericidal function. Significantly elevated activation levels of Neu_CSTA and Neu_classic subsets were observed in *Fpr2*-KO compared to WT mice, indicating that *Fpr2* deletion may result in certain dysregulations which may be very unfavorable for PMN to play a bactericidal role in the infected state.

The examination of inter-subpopulation regulation revealed that the Neu_CCL4 subset predominantly functions as a regulator in immune homeostasis. Despite its limited size, this subset exerts a significant regulatory influence on other subsets. The absence of the Neu_CCL4 cluster disrupts PMN sterilization-related functions within homeostasis. Conversely, during infection, the regulatory impact of Neu_CCL4 diminishes, leading to enhanced interactions among various subsets and ultimately improving the bactericidal efficacy of PMN. The analysis indicates a potential relationship between the regulatory homeostasis of Neu_CCL4 and the expression of the ANXA1 pathway, which possesses anti-inflammatory properties via Fpr2 (Ni et al., 2021; Perretti & D’Acquisto, 2009). This suggests that the regulatory impact of Neu_CCL4 on PMN homeostasis may be modulated by ANXA1, with Fpr2 potentially acting as a crucial checkpoint. *ANXA1* expression in Neu_CCL4 exhibited a statistically significant correlation with ELF4 regulon in the rest condition.

ELF4 (E74 like ETS transcription factor 4) is an X-linked ETS transcription factor, which is highly expressed in hematopoietic cells and regulates many kinds of cells from hematopoietic stem cells to peripheral blood immune cells and plays an important role in regulating host immunity (Suico, Shuto, & Kai, 2017). ELF4 plays a crucial role in modulating the host’s inflammatory response as a significant transcriptional regulator that serves to restrict excessive inflammation. Its functional deficiency will lead to fever and ulcers in the gastrointestinal tract, especially in the oral cavity, which is related to the increased expression of innate cytokines (Tyler et al., 2021). Mucosal-related inflammation in patients with clinical Elf4-X linkage deficiency(DEX) is aggravated, which is characterized by neutrophil infiltration, increased expression of IL-17, and increased pro-inflammatory response of macrophages. Macrophages lacking Elf4 showed inflammatory hyperreactivity and the expression of anti-inflammatory genes was defective and the expression of pro-inflammatory genes was enhanced (Curina et al., 2017; Tyler et al., 2021). Studies have shown that for LPS-stimulated BMDM, the anti-inflammatory gene IL1RN needs the presence of Elf4 to be continuously expressed(Tyler et al., 2021). Research has demonstrated that ELF4 can suppress the differentiation of Th17 cells. The absence of Elf4 in mice results in a notable elevation in IL-17 levels, which has been linked to the anti-infective properties of PMN (Lee et al., 2014). *Fpr2* is highly expressed in myeloid cells. In this study, the deletion of *Fpr2* leads to the up-regulation of inflammation-related factors. Although it is not as serious as the deletion of *Elf4*, the expression of IL-17, IL-6, and *S100a8*/*S100a9* are significantly up-regulated with the latter being a potent pro-inflammatory factor. Previous studies have suggested that Elf4 controls the homeostasis, activation, and homing of T cells. Here, we propose that this gene may control the homeostasis of PMN by regulating the expression of *Anxa1*.

According to previous studies and databases, there is no report about the direct combination of Elf4 and *Anxa1*. However, some studies showed that as a transcription regulator, the regulatory effect of Elf4 on CD8+T cells is at least partly caused by the direct regulation of the *Klf4* gene (Yamada et al., 2009). And Klf4 can directly regulate the expression of *Anxa1* (Chen et al., 2023). Therefore, we speculate that Elf4, which is specifically expressed in Neu_CCL4 at steady state, may regulate *Anxa1* expression through Klf4. In the infection state, it shows that the expressions of *Elf4*, *Klf4*, and *Anxa1* are significantly down-regulated, which is consistent with our conjecture, that is, these three genes are important regulatory genes for PMN to maintain a stable immune balance. However, the specific mechanism of interaction between Elf4 and Klf4 and the mechanism of regulating *Anxa1* expression needs to be further proved. Furthermore, our research has revealed that the modulation of Elf4 and *Anxa1* expression is influenced by Fpr2, suggesting the presence of a potential positive feedback regulatory pathway involving these genes.

Besides regulating inflammation, Elf4 can also regulate the development of NK cells and the expression of perforin. The expression of type I IFN in NK cells lacking *Elf4* is down-regulated, resulting in the lack of antiviral and antitumor effects of NK and NKT cells (You et al., 2013). At the same time, Elf4 is induced by type I interferon and its feedback stimulates the production of interferon and the expression of interferon stimulating gene. However, in this study, we found that when the infection occurred, the expression of *Elf4* in both mice decreased significantly, but the expression of type I IFN in KO mice increased significantly, indicating that the expression of type I IFN may have little to do with Elf4 in the infected state, which further confirmed that the gene may be more important in regulating the steady state of PMN.

Further analysis showed that the pathway of type I and II interferons increased significantly in Neu_CCL4 and Neu_ISG clusters, with Neu_CCL4 being the most significant. It is known that type I IFN has anti-inflammatory function, especially through negative regulation of IL-1 and IL-18 and inflammatory Th17 cells (Guarda et al., 2011; Guo, Chang, & Cheng, 2008; Reboldi et al., 2014). It has also been reported that IFN-γ secreted by NK cells can promote the function of PMN by inhibiting its apoptosis and promoting it to retain its bactericidal function (Bhatnagar et al., 2010). Here, whether there is a correlation between the down-regulation of bactericidal function and IFN expression in KO mice needs further study.

At the time of infection, we found that the communication between clusters was obviously strengthened and the transcription of anti-infection related genes in each group was significantly increased, indicating that different functional subgroups may cooperate to finally realize the anti-infection function. However, the significant difference in anti-infection gene expression between WT and KO mice is still concentrated in the Neu_classic and Neu_CSTA subgroups, indicating that these two subgroups are still the main functional groups.

In conclusion, we proposed for the first time that Neu_CCL4 cluster in peripheral blood PMN may play a function similar to Treg in the steady state of PMN, and Elf4 regulon in this cluster plays an important regulatory role, which is related to Fpr2. The expression of *Anxa1* is regulated by Elf4 and Anxa1 exerts its immune balance regulation function by binding to Fpr2 of each cluster. However, when infection occurred, the expression of *Elf4* was significantly down-regulated and the corresponding expression of *Anxa1* was also down-regulated, so that the inhibition of Neu_CCL4 was down-regulated, the interaction among cluster was enhanced and the bactericidal function of PMN was increased.

## Materials and Methods

### Single-cell transcriptomic analysis

The peripheral blood was taken into an EDTA anticoagulation tube. The red blood cells were lysed on ice for 3min and centrifuged at 2,300 rpm for 3 min, while the supernatant was removed to obtain the cells. Fastq-format files were mapped to the mouse genome(mm10 from 10x genomics) by CellRanger software(version: 5.0.1). Then we employed the Seurat package(version: 4.9.9.9045) in R platform(version: 4.2.1) for quality control, gene raw count normalization, variable features selection and sample integration (Hao et al., 2021). In order to filter out cells with low quality, cells with the gene number less than 200 or count less than 2000 were removed. Mitochondrial genes were defined as gene names beginning with “mt-” and cells with high mitochondrial gene ratio(≥10%) were also depleted. Cells with gene number greater than 5000 were seemed as doublets and removed from subsequent analysis. Dataset of each sample was normalized and scaled by “SCTransform” function (Hafemeister & Satija, 2019) and further processed with “integration” to remove the batch effect (Stuart et al., 2019). Principal component analysis(PCA) was conducted based on the scaled data with the top 2000 high variable genes and the top 30 principals were used for Uniform Manifold Approximation and Projection(UMAP) analysis. Using the graph-based cluster method, we acquired the unsupervised cell cluster results by FindNeighbors and FindClusters function. Then these cell clusters were annotated according to their canonical markers based on the previous research, that is Ptprc for all immune cells, *Cd68*/*CD300e*/*Apoe* for macrophages, *Csf3r/Il1b*/*Cxcr2* for neutrophils, *H2-DMa*/*H2-DMb1* for dendritic cells(DCs), *Ly6c2*/*Ccr2*/*Fcgr1* for monocytes. Cells expressed markers of different lineages(*i.e.*, T lineage and B lineage) were also defined as doublets and removed. For subcluster analysis, cells in a specific cell cluster were selected to repeat the procedures including PCA, UMAP, markers identification and cluster annotation with the same variable features as total cells.

### Differentially expressed gene(DEG) and enrichment analysis

To identify differentially expressed genes among samples, the function “FindMarkers” with Wilcox rank-sum test algorithm was applied with the following parameters: min.pct = 0.4, min.diff.pct = 0.05, logfc.threshold = 0.25. The list of interested genes for enrichment analysis was passed to WebGestaltR package(version: 0.4.6) with following parameters: enrichMethod = "ORA", organism = "hsapiens", fdrThr = 0.2, referenceSet = "genome", enrichDatabase = "pathway_KEGG". Then the results were shown by bar plot. Unique pathways were defined by following criteria: FDR < 0.05, gene num > 3, duplicated pathways in each cell type or pathways shared same genes with other pathways but had lower enrichment ratio were removed. For single-cell gene-set enrichment analysis(GSEA), SCT matrix from Seurat object was passed to gsva function(method="ssgsea", ssgsea.norm=T) in GSVA package(version: 1.44.5). Hallmarks gene set, which was downloaded through msigdbr package(version: 7.5.1), was chosen as the reference.

### Pseudo-time analysis

Pseudo-time analysis was conducted by monocle3(version: 1.2.9) for developmental trajectories inference (Cao et al., 2019; Qiu et al., 2017; Trapnell et al., 2014). Raw counts matrix and meta data from the integrated Seurat object were passed to monocle3 workflow to form a monocle3 cell dataset(cds) object. Firstly, the cds object was preprocessed to prepare for trajectory inference by “preprocess_cds” function. Then the UMAP embedding calculated by “reduce_dimension” function was replaced by the corresponding values stored in Seurat object. Subsequently, cells were clustered using “cluster_cells” function and the principal graph was learned from the reduced dimension space using the “learn_graph” function with default parameters. Finally, pseudotime score were calculated using the “order_cells” function and the trajectory was visualized using the “plot_cells” function.

### Cell-cell communication analysis

CellChat(version 1.5.0, www.cellchat.org) is a publicly available repository of curated receptors, ligands and their interactions, that can be used to search for cell-cell interaction and receptor-ligand pairs among cell types (Jin et al., 2021). Potential interactions between the two cell types were inferred by using the CellChat method based on expression quantification levels of receptor and ligand gene pairs. Only receptors and ligands expressed in more than 10% of the cells in the corresponding subclusters were considered. We compared the signaling network similarity between *Fpr2*-KO and WT in *GBS* uninfected or infected situation respectively.

### Regulon module prediction through SCENIC

Single-Cell rEgulatory Network Inference and Clustering(SCENIC) analysis was performed to reveal the gene regulatory network(GRN) in different cell types and clusters. We performed the SCENIC analysis using the latest version of pySCENIC(version: 0.10.2), a lightning-fast python implementation of the SCENIC pipeline. The gene-motif rankings(10kb upstream and downstream of the transcription start site[TSS]) were used to determine the search space around the TSS. The motif database(mc9nr) including 24,453 motifs was used for RcisTarget and GENIE3 algorithms to infer the GRNs, respectively. Regulon specificity score(RSS) was calculated by calcRSS function of SCENIC package(version: 1.3.1) in R.

### Blood sampling handling and neutrophil isolation

#### PMNs isolation

4mL blood was layered on top of 4 mL of 1113 ( 1113, Invitrogen, USA) and 3mL 1115 (1115, Invitrogen, USA) in a 15mL centrifugation tube. The tube was centrifuged at 800g at 20°C for 30min. The PMN cell layer was collected and red blood cells were removed using Red Blood Lysing Buffer (555899, BD Pharm Lysing Buffer).

#### Neu_CSTA isolation

Freshly isolated neutrophils were suspended at 1×10^7^/mL in PBS. For separation of CSTA^+^ neutrophils, cells labeled with MACS beads are captured by the magnetic field of the separator (130-090-312, Miltenyi Biotec Inc.), whereas unlabeled cells pass the magnetic field and end up in the flow through fraction. In short, freshly isolated neutrophils were resuspended in 100μL MACS Separation Buffer (130-091-221, Miltenyi Biotec Inc.) and stained with APC-conjugated anti-human CSTA antibody (2μL/10^6^ cells) for 10min in the dark and cells were washed twice. Then incubated with anti-APC microbeads (130-100-070, 20μL/10^7^ cells, Miltenyi Biotec Inc.) in 80μL MACS Separation Buffer (130-091-221, Miltenyi Biotec Inc.) for 15min in the dark. Add 1-2 mL of buffer to wash the cells and centrifuge at 300g for 10min. Subsequently, the CSTA^+^ neutrophils were separated on an MS column (130-042-201, Miltenyi Biotec Inc.) on a MACS Separator (130-090-312, Miltenyi Biotec Inc.) and washed again to detach from the antibody-magnetic bead.

#### Neu_CCL4 isolation

For separation of CCL4^+^ neutrophils, cells labeled with MACS beads are captured by the magnetic field of the separator (130-090-312, Miltenyi Biotec Inc), whereas unlabeled cells pass the magnetic field and end up in the flow through fraction. In short, freshly isolated neutrophils were resuspended in 100μL MACS Separation Buffer (130-091-221, Miltenyi Biotec Inc.) and stained with PE-conjugated anti-human CCL4 antibody (2 μL/10^6^ cells) for 10min in the dark and cells were washed twice. Then incubated with anti-PE microbeads (2130-048-80, 10μL/10^7^ cells, Miltenyi Biotec Inc.) in 80μL MACS Separation Buffer (130-091-221, Miltenyi Biotec Inc.) for 15 min in the dark. Add 1-2mL of buffer to wash the cells and centrifuge at 300g for 10min. Subsequently, the CCL4^+^ neutrophils were separated on an MS column (130-042-201, Miltenyi Biotec Inc.) on a MACS Separator (130-090-312, Miltenyi Biotec Inc.) and washed again to detach from the antibody-magnetic bead. On the other hand, the CCL4^-^ neutrophils were separated on an LD column (130-042-001, Miltenyi Biotec Inc.) on a MiniMACS Separator (130-042-102, Miltenyi Biotec Inc.) and washed again to detach from the antibody-magnetic bead. The isolated neutrophils were resuspended in PBS for subsequent experiments. The purity of the cells was routinely 90 to 95%.

### ROS detection

For Neu_CSTA cluster ROS analysis, 5 × 10^5^ neutrophils were isolated from human and mouse and treated with Flow Cytometry antibodies at 37°C for 30 min, then cells stained with FITC anti-human CD15 (301904, BioLegend, Trustain, USA) and APC anti-mouse/human CSTA and intracellular ROS production was measured by a dihydroethidium probe kit (BB-47051, DHE, BestBio, Beijing, China) at 37°C for 30min. After washing, the cells were immediately analyzed by flow cytometry. For PMN ROS analysis, we use qPCR to detect relative gene expression (**Supplementary Table S3**).

### Flow Cytometry analysis

After the mice were infected with *GBS* for 3 hours, the peripheral blood was taken into an EDTA anticoagulation tube. The red blood cell was lysed on ice for three minutes and centrifuged at 2,300rpm for 3min and then the supernatant was removed to obtain the cells. Cells were resuspended in FACS buffer (BD Bioscience) and incubated with Fc blocking antibody (BioLegend, Trustain, USA) for 10min to prevent nonspecific binding. The absolute count of blood leukocytes in the blood and peritoneal lavage fluid samples were determined using a BD TruCount system (BD Biosciences, USA). The cells were stained with PerCP/Cyanine5.5 anti-mouse Ly6G, APC-Cy7 anti-mouse CD45, FITC-anti-mouse CD11b, FITC anti-human CD15 and PE anti-human CD63, PerCP/Cyanine5.5 anti-human CD63. All antibodies were purchased from Biolegend (Trustain, USA). The cells were stained with APC anti-mouse/human CSTA, PE anti-mouse/human CCL4 (Univ, China). All of the above-stained cells were assayed using a BD FACSCelesta™ flow cytometer and data were analyzed using FlowJo software.

### Statistical analysis

The specific tests used to analyze each set of experiments are indicated in the Figure legends. The GraphPad Prism software (GraphPad Software, San Diego, California) and R software (https://www.r-project.org/) were employed for the statistical calculations and Figure visualization. Comparisons between two groups were performed using a two-tailed unpaired Student’s t-test. \**p* < 0.05, \*\**p* <0.01. ns, denotes not significant.

### Ethics statement

This research was performed in compliance with the guidelines of laboratory animal care and was approved in China. All experimental procedures were approved by the Animal Ethics Committee of the Academy of Military Medical Sciences. The use of samples from all sources has been approved by the Ethics Committee of Beijing Dongfang Hospital and all participants have signed informed consent forms. Ethical approval number: JDF-IRB-2020041002.

## Data Availability

All single-cell sequencing data have been deposited at the National Genomics Data Center with accession code PRJCA026504. The data analysis codes and any additional information are available from the corresponding author upon reasonable request.

## Acknowledgments

This work was supported by the State Key Laboratory of Pathogen and Biosecurity, No.SKLPBS2212.

## Author Contributions

Z.S. and S.M. performed the whole experiment and analysis. Z.S. wrote the first draft of the manuscript. H.J. and P.Z. designed the study and revised the manuscript. All authors contributed to the manuscript revision and read and approved the submitted version.

## References

Ahmed S., O. A. Odumade, P. van Zalm, B. Fatou, R. Hansen, C. R. Martin, A. Angelidou, & H. Steen. 2023. Proteomics-Based Mapping of Bronchopulmonary Dysplasia-Associated Changes in Noninvasively Accessible Oral Secretions. J Pediatr, 270: 113774. doi:10.1016/j.jpeds.2023.113774,

Beyrau M., J. V. Bodkin, & S. Nourshargh. 2012. Neutrophil heterogeneity in health and disease: a revitalized avenue in inflammation and immunity. Open Biol, 2: 120134. doi:10.1098/rsob.120134,

Bhatnagar N., H. S. Hong, J. K. Krishnaswamy, A. Haghikia, G. M. Behrens, R. E. Schmidt, & R. Jacobs. 2010. Cytokine-activated NK cells inhibit PMN apoptosis and preserve their functional capacity. Blood, 116: 1308–1316. doi:10.1182/blood-2010-01-264903,

Cao J., M. Spielmann, X. Qiu, X. Huang, D. M. Ibrahim, A. J. Hill, F. Zhang, S. Mundlos, L. Christiansen, F. J. Steemers, C. Trapnell, & J. Shendure. 2019. The single-cell transcriptional landscape of mammalian organogenesis. Nature, 566: 496–502. doi:10.1038/s41586-019-0969-x,

Chen Y., S. Zhu, T. Liu, S. Zhang, J. Lu, W. Fan, L. Lin, T. Xiang, J. Yang, X. Zhao, Y. Xi, Y. Ma, G. Cheng, D. Lin, & C. Wu. 2023. Epithelial cells activate fibroblasts to promote esophageal cancer development. Cancer Cell, 41: 903–918 e908. doi:10.1016/j.ccell.2023.03.001,

Consortium E. P. 2012. An integrated encyclopedia of DNA elements in the human genome. Nature, 489: 57–74. doi:10.1038/nature11247,

Corminboeuf O., & X. Leroy. 2015. FPR2/ALXR agonists and the resolution of inflammation. J Med Chem, 58: 537–559. doi:10.1021/jm501051x,

Curina A., A. Termanini, I. Barozzi, E. Prosperini, M. Simonatto, S. Polletti, A. Silvola, M. Soldi, L. Austenaa, T. Bonaldi, S. Ghisletti, & G. Natoli. 2017. High constitutive activity of a broad panel of housekeeping and tissue-specific cis-regulatory elements depends on a subset of ETS proteins. Genes Dev, 31: 399–412. doi:10.1101/gad.293134.116,

Deerhake M. E., E. Y. Reyes, S. Xu-Vanpala, & M. L. Shinohara. 2021. Single-Cell Transcriptional Heterogeneity of Neutrophils During Acute Pulmonary Cryptococcus neoformans Infection. Front Immunol, 12: 670574. doi:10.3389/fimmu.2021.670574,

Devchand P. R., M. Arita, S. Hong, G. Bannenberg, R. L. Moussignac, K. Gronert, & C. N. Serhan. 2003. Human ALX receptor regulates neutrophil recruitment in transgenic mice: roles in inflammation and host defense. FASEB J, 17: 652–659. doi:10.1096/fj.02-0770com,

Dufton N., R. Hannon, V. Brancaleone, J. Dalli, H. B. Patel, M. Gray, F. D’Acquisto, J. C. Buckingham, M. Perretti, & R. J. Flower. 2010. Anti-inflammatory role of the murine formyl peptide receptor 2: ligand-specific effects on leukocyte responses and experimental inflammation. J Immunol, 184: 2611–2619. doi:10.4049/jimmunol.0903526,

Guarda G., M. Braun, F. Staehli, A. Tardivel, C. Mattmann, I. Forster, M. Farlik, T. Decker, R. A. Du Pasquier, P. Romero, & J. Tschopp. 2011. Type I interferon inhibits interleukin-1 production and inflammasome activation. Immunity, 34: 213–223. doi:10.1016/j.immuni.2011.02.006,

Guo B., E. Y. Chang, & G. Cheng. 2008. The type I IFN induction pathway constrains Th17-mediated autoimmune inflammation in mice. J Clin Invest, 118: 1680–1690. doi:10.1172/JCI33342,

Hafemeister C., & R. Satija. 2019. Normalization and variance stabilization of single-cell RNA-seq data using regularized negative binomial regression. Genome Biol, 20: 296. doi:10.1186/s13059-019-1874-1,

Hao Y., S. Hao, E. Andersen-Nissen, W. M. Mauck, 3rd, S. Zheng, A. Butler, M. J. Lee, A. J. Wilk, C. Darby, M. Zager, P. Hoffman, M. Stoeckius, E. Papalexi, E. P. Mimitou, J. Jain, A. Srivastava, T. Stuart, L. M. Fleming, B. Yeung, A. J. Rogers, J. M. McElrath, C. A. Blish, R. Gottardo, P. Smibert, & R. Satija. 2021. Integrated analysis of multimodal single-cell data. Cell, 184: 3573-3587 e3529. doi:10.1016/j.cell.2021.04.048,

He H. Q., & R. D. Ye. 2017. The Formyl Peptide Receptors: Diversity of Ligands and Mechanism for Recognition.Molecules, 22. doi:10.3390/molecules22030455,

Jin S., C. F. Guerrero-Juarez, L. Zhang, I. Chang, R. Ramos, C. H. Kuan, P. Myung, M. V. Plikus, & Q. Nie. 2021. Inference and analysis of cell-cell communication using CellChat. Nat Commun, 12: 1088. doi:10.1038/s41467-021-21246-9,

Kolaczkowska E., & P. Kubes. 2013. Neutrophil recruitment and function in health and inflammation. Nat Rev Immunol, 13: 159–175. doi:10.1038/nri3399,

Kretschmer D., A. K. Gleske, M. Rautenberg, R. Wang, M. Koberle, E. Bohn, T. Schoneberg, M. J. Rabiet, F. Boulay, S. J. Klebanoff, K. A. van Kessel, J. A. van Strijp, M. Otto, & A. Peschel. 2010. Human formyl peptide receptor 2 senses highly pathogenic Staphylococcus aureus. Cell Host Microbe, 7: 463–473. doi:10.1016/j.chom.2010.05.012,

Lee P. H., M. Puppi, K. S. Schluns, L. Y. Yu-Lee, C. Dong, & H. D. Lacorazza. 2014. The transcription factor E74-like factor 4 suppresses differentiation of proliferating CD4+ T cells to the Th17 lineage. J Immunol, 192: 178–188. doi:10.4049/jimmunol.1301372,

Li L., Y. Pian, S. Chen, H. Hao, Y. Zheng, L. Zhu, B. Xu, K. Liu, M. Li, H. Jiang, & Y. Jiang. 2016. Phenol-soluble modulin alpha4 mediates Staphylococcus aureus-associated vascular leakage by stimulating heparin-binding protein release from neutrophils. Sci Rep, 6: 29373. doi:10.1038/srep29373,

Liu M., K. Chen, T. Yoshimura, Y. Liu, W. Gong, A. Wang, J. L. Gao, P. M. Murphy, & J. M. Wang. 2012. Formylpeptide receptors are critical for rapid neutrophil mobilization in host defense against Listeria monocytogenes. Sci Rep, 2: 786. doi:10.1038/srep00786,

Machado M. G., L. P. Tavares, G. V. S. Souza, C. M. Queiroz-Junior, F. R. Ascencao, M. E. Lopes, C. C. Garcia, G. B. Menezes, M. Perretti, R. C. Russo, M. M. Teixeira, & L. P. Sousa. 2020. The Annexin A1/FPR2 pathway controls the inflammatory response and bacterial dissemination in experimental pneumococcal pneumonia. FASEB J, 34: 2749–2764. doi:10.1096/fj.201902172R,

Ni C., S. Gao, X. Li, Y. Zheng, H. Jiang, P. Liu, Q. Lv, W. Huang, Q. Li, Y. Ren, Z. Mi, D. Kong, & Y. Jiang. 2023. Fpr2 exacerbates Streptococcus suis-induced streptococcal toxic shock-like syndrome via attenuation of neutrophil recruitment. Front Immunol, 14: 1094331. doi:10.3389/fimmu.2023.1094331,

Ni C., S. Gao, Y. Zheng, P. Liu, Y. Zhai, W. Huang, H. Jiang, Q. Lv, D. Kong, & Y. Jiang. 2021. Annexin A1 Attenuates Neutrophil Migration and IL-6 Expression through Fpr2 in a Mouse Model of Streptococcus suis-Induced Meningitis. Infect Immun, 89. doi:10.1128/IAI.00680-20,

Perretti M., & F. D’Acquisto. 2009. Annexin A1 and glucocorticoids as effectors of the resolution of inflammation. Nat Rev Immunol, 9: 62–70. doi:10.1038/nri2470,

Qi X., Y. Yu, R. Sun, J. Huang, L. Liu, Y. Yang, T. Rui, & B. Sun. 2021. Identification and characterization of neutrophil heterogeneity in sepsis. Crit Care, 25: 50. doi:10.1186/s13054-021-03481-0,

Qiu X., Q. Mao, Y. Tang, L. Wang, R. Chawla, H. A. Pliner, & C. Trapnell. 2017. Reversed graph embedding resolves complex single-cell trajectories. Nat Methods, 14: 979–982. doi:10.1038/nmeth.4402,

Reboldi A., E. V. Dang, J. G. McDonald, G. Liang, D. W. Russell, & J. G. Cyster. 2014. Inflammation. 25-Hydroxycholesterol suppresses interleukin-1-driven inflammation downstream of type I interferon. Science, 345: 679-684. doi:10.1126/science.1254790,

Seifert R., & K. Wenzel-Seifert. 2001. Defective Gi protein coupling in two formyl peptide receptor mutants associated with localized juvenile periodontitis. J Biol Chem, 276: 42043–42049. doi:10.1074/jbc.M106621200,

Silvestre-Roig C., A. Hidalgo, & O. Soehnlein. 2016. Neutrophil heterogeneity: implications for homeostasis and pathogenesis. Blood, 127: 2173–2181. doi:10.1182/blood-2016-01-688887,

Simiele F., A. Recchiuti, D. Mattoscio, A. De Luca, E. Cianci, S. Franchi, V. Gatta, A. Parolari, J. P. Werba, M. Camera, B. Favaloro, & M. Romano. 2012. Transcriptional regulation of the human FPR2/ALX gene: evidence of a heritable genetic variant that impairs promoter activity. FASEB J, 26: 1323–1333. doi:10.1096/fj.11-198069,

Stuart T., A. Butler, P. Hoffman, C. Hafemeister, E. Papalexi, W. M. Mauck, 3rd, Y. Hao, M. Stoeckius, P. Smibert, & R. Satija. 2019. Comprehensive Integration of Single-Cell Data. Cell, 177: 1888-1902 e1821. doi:10.1016/j.cell.2019.05.031,

Suico M. A., T. Shuto, & H. Kai. 2017. Roles and regulations of the ETS transcription factor ELF4/MEF. J Mol Cell Biol, 9: 168–177. doi:10.1093/jmcb/mjw051,

Sun Z., W. Huang, Y. Zheng, P. Liu, W. Yang, Z. Guo, D. Kong, Q. Lv, X. Zhou, Z. Du, H. Jiang, & Y. Jiang. 2021. Fpr2/CXCL1/2 Controls Rapid Neutrophil Infiltration to Inhibit Streptococcus agalactiae Infection. Front Immunol, 12: 786602. doi:10.3389/fimmu.2021.786602,

Trapnell C., D. Cacchiarelli, J. Grimsby, P. Pokharel, S. Li, M. Morse, N. J. Lennon, K. J. Livak, T. S. Mikkelsen, & J. L. Rinn. 2014. The dynamics and regulators of cell fate decisions are revealed by pseudotemporal ordering of single cells. Nat Biotechnol, 32: 381–386. doi:10.1038/nbt.2859,

Tyler P. M., M. L. Bucklin, M. Zhao, T. J. Maher, A. J. Rice, W. Ji, N. Warner, J. Pan, R. Morotti, P. McCarthy, A. Griffiths, A. M. C. van Rossum, I. Hollink, V. Dalm, J. Catanzaro, S. A. Lakhani, A. M. Muise, & C. L. Lucas. 2021. Human autoinflammatory disease reveals ELF4 as a transcriptional regulator of inflammation. Nat Immunol, 22: 1118–1126. doi:10.1038/s41590-021-00984-4,

Weiss E., & D. Kretschmer. 2018. Formyl-Peptide Receptors in Infection, Inflammation, and Cancer. Trends Immunol, 39: 815–829. doi:10.1016/j.it.2018.08.005,

Xie X., Q. Shi, P. Wu, X. Zhang, H. Kambara, J. Su, H. Yu, S. Y. Park, R. Guo, Q. Ren, S. Zhang, Y. Xu, L. E. Silberstein, T. Cheng, F. Ma, C. Li, & H. R. Luo. 2020. Single-cell transcriptome profiling reveals neutrophil heterogeneity in homeostasis and infection. Nat Immunol, 21: 1119–1133. doi:10.1038/s41590-020-0736-z,

Yamada T., C. S. Park, M. Mamonkin, & H. D. Lacorazza. 2009. Transcription factor ELF4 controls the proliferation and homing of CD8+ T cells via the Kruppel-like factors KLF4 and KLF2. Nat Immunol, 10: 618–626. doi:10.1038/ni.1730,

You F., P. Wang, L. Yang, G. Yang, Y. O. Zhao, F. Qian, W. Walker, R. Sutton, R. Montgomery, R. Lin, A. Iwasaki, & E. Fikrig. 2013. ELF4 is critical for induction of type I interferon and the host antiviral response. Nat Immunol, 14: 1237–1246. doi:10.1038/ni.2756,

Zhang H., Y. Lu, G. Sun, F. Teng, N. Luo, J. Jiang, & A. Wen. 2017. The common promoter polymorphism rs11666254 downregulates FPR2/ALX expression and increases risk of sepsis in patients with severe trauma. Crit Care, 21: 171. doi:10.1186/s13054-017-1757-3,

Zhang M., J. L. Gao, K. Chen, T. Yoshimura, W. Liang, W. Gong, X. Li, J. Huang, D. H. McDermott, P. M. Murphy, X. Wang, & J. M. Wang. 2020. A Critical Role of Formyl Peptide Receptors in Host Defense against Escherichia coli. J Immunol, 204: 2464–2473. doi:10.4049/jimmunol.1900430,

